# FoldToken4: Consistent & Hierarchical Fold Language

**DOI:** 10.1101/2024.08.04.606514

**Authors:** Zhangyang Gao, Cheng Tan, Stan Z. Li

## Abstract

Creating protein structure language has attracted increasing attention in unifing the modality of protein sequence and structure. While recent works, such as FoldToken1&2&3 have made great progress in this direction, the relationship between languages created by different models at different scales is still unclear. Moreover, models at multiple scales (different code space size, like 2^5^, 2^6^, ⋯, 2^12^) need to be trained separately, leading to redundant efforts. We raise the question: *Could a single model create multiscale fold languages?* In this paper, we propose FoldToken4 to learn the consistent and hierarchical of multiscale fold languages. By introducing multiscale code adapters and token mixing techniques, FoldToken4 can generate multiscale languages from the same model, and discover the hierarchical token-mapping relationships across scales. To the best of our knowledge, FoldToken4 is the first effort to learn multi-scale token consistency and hierarchy in VQ research; Also, it should be more novel in protein structure language learning.

## 1 Introduction

“SE-(3) structure should not be special and difficult. Let’s lower the barrier.”

– Our Goal

Creating protein structure language has attracted increasing attention in unifing the modeling paradigm of sequence and structure. Recent works, such as FoldToken1, FoldToken2, and FoldToken3, have made great progress in this direction: FoldToken2 improves the reconstruction RMSD by 80% compared to FoldToken1, and FoldToken3 further enhances the compression ratio using 0.39% code space of FoldToken2. Generally, there is a balance between compression and reconstruction: model with smaller code space captures the coarse geometry with higher compression ability, and vise versa. The multi-scale consistency and hierarchy of fold language would be valuable for understanding and analyzing protein structures. Unfortunately, current vector quantization works have ignored this. *How to associate multiscale languages and discover their intristic laws remain an open question*.

Existing fold tokenization methods are limited to single-scale expression ability. For example, FoldToken2 [7], ESM3 [9], and FoldToken3 [3] have the codebook size of 65536, 4096, and 256, respectively. By gradually reducing the codebook size, one can get a better compression ratio, while the reconstruction quality is compromised, resulting in a coarse-grained hierarchical representation. The consistency and hierarchy of fold language across different scales remain unknown, as models at different scales are trained separately, lacking a unified representation space for knowledge sharing. Consequently, there is no way to analyze the language’s consistency and hierarchy without building another model to translate tokens across scales. In addition, the training of models at different scales is time-consuming and resource-intensive, and saving codes for each scale violates the compression principle. *Can we design one model to generate semantically consistent multiscale tokens without redundance in both model training and code saving?*

We propose FoldToken4 to create a consistent and hierarchical fold language across scales. Firstly, we make multi-scale models share the same encoder to learn a **consistent** semantic space. To align multi-scale tokens with encoder’s output, we introduce code adapters to generate code embeddings at different scales and mix codes across scales. Secondly, we can compute the token embedding similarity to reveal translation relationships across scales, recorded by the transition matrix that reflects the **hierarchy** of the fold language. This hierarchy helps to understand and visualize protein structures in a coarse-to-fine manner. Finally, FoldToken4 can remove redundant efforts in training and code saving: One just needs to train one model for all scales, and save the finest-scale code, which can be translated to any other scale using the transition matrix without rerunning the model.

We evaluate FoldToken4 on both single-chain and multi-chain reconstruction tasks, showing that it achieves competitive but slightly worse reconstruction quality than FoldToken3. However, FoldToken4 further compresses the code space to 32, and multi-scale languages, such as 32, 64, 128, 256, ⋯, 4096, can be generated from the same model. In summary, FoldToken4 offers advantages in consistency, hierarchy, and efficiency, in addition to FoldToken2&3’s invariant, compact, and generative merits. We believe FoldToken4 will be valuable for analyzing fold languages from a coarse-to-fine perspective and benefit a wide range of protein structure-related tasks.

## 2 Method

### 2.1 Overall Framework

As shown in Fig.1, the overall framework keeps the same as FoldToken1 [5, 7], FoldToken2 [6] and FoldToken3 [3], including encoder, quantifier and decoder:

**Figure 1:**
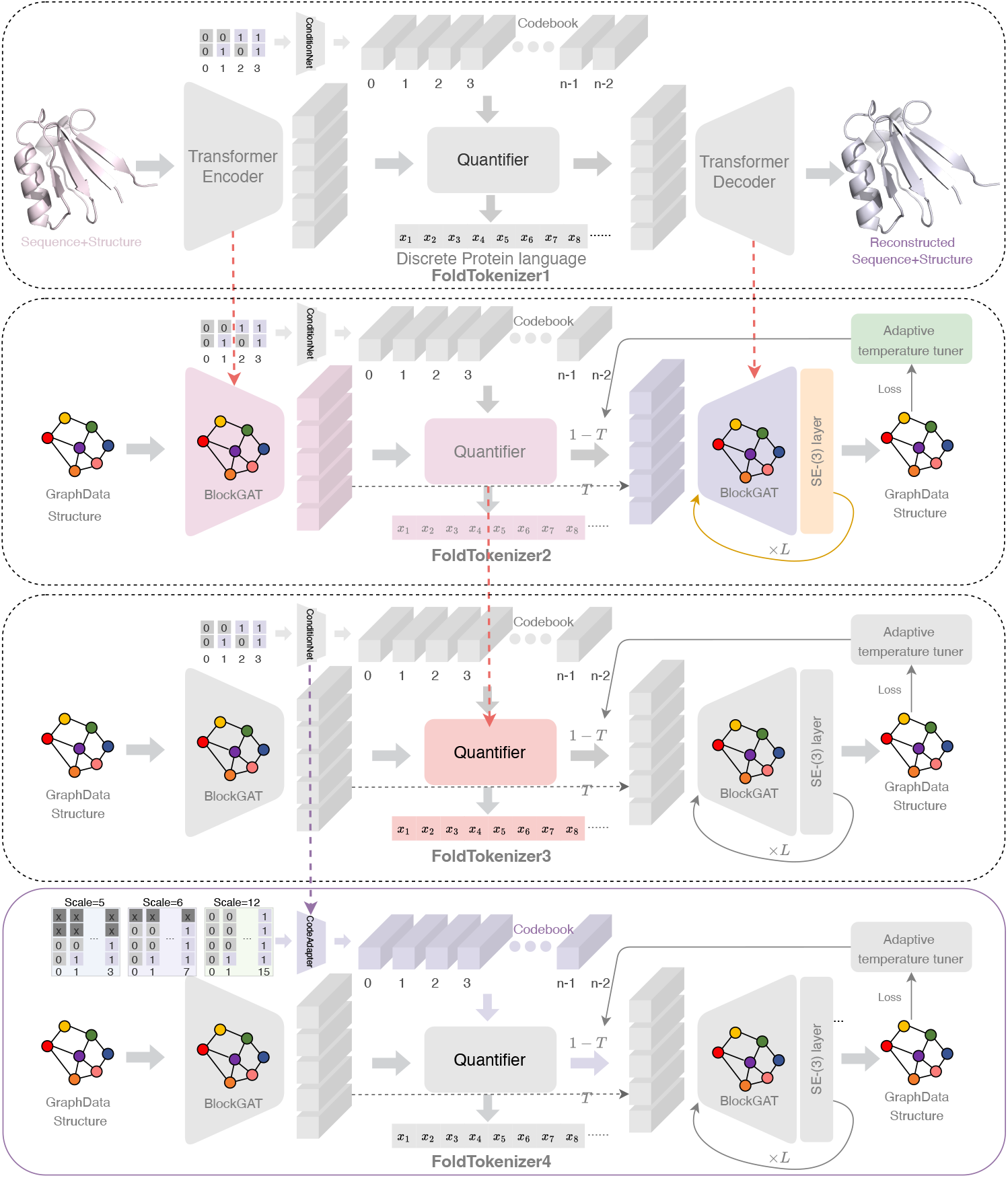
The overall framework of FoldTokenizer4, which contains contains encoder, quantifier, and decoder. We use BlockGAT to encoder protein structures as invariant embeddings, BSQ to quantize the embeddings into discrete tokens, and SE-(3) layer to recover the protein structures iteratively.

**Brief Overview**

**Step1: Encoding**.Given the protein *𝒢*, we use BlockGAT [8] to learn residue features:

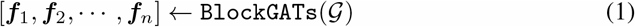

where *f* is the learned features of the s-th residue.

**Step2: Quantization**. Given **F** = [*f*_1_, *f*_2_, ⋯, *f*_n_], we quantize the embeddings into discrete tokens (**VQ-IDs**) Z = [z_1_, z_2_, ⋯, z_n_] using the binary stochastic quantifier (**BSQ**). The quantifier (Q : ***f*** ↦ *z*) and de-quantifier 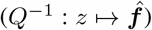 converts continuous embedding ***f*** as a discrete latent code *z*, and vise versa:

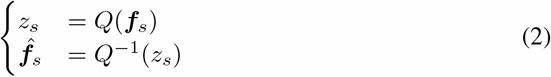

By setting the codebook size as 2^m^, the discrete tokens are restricted to [0, ^2m™1^]. Similar to 20 amino acid types that describing protein sequence, VQ-IDs are used to represent the protein structures. The quantized sequence Z = [z_1_, z_2_, ⋯, z_n_] is termed as fold fanguage.

**Step3: Decoding**. We apply the SE-(3) BlockGAT Decoder to recover the protein structures:

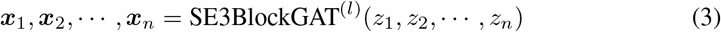

where x_s_ ∈ ℝ^4,3^ is the coordinates of the s-th residue’s backbone atoms.

#### Contribution

From FoldToken3 to FoldToken4, we make the following improvements:

1. **Scale-aware Code Adapter**: We introduce code adapters to generate code embeddings at different scales, all of whcih are aligned with the encoder’s output embedding space.
2. **Multiscale Mix-Token Training**: Multiscale models, with code book size of 2^5^, 2^6^, ⋯, 2^12^, are trained simultaneously, avoiding redundant efforts in model training. The shared encoder, quantifier, and decoder also provide knowledge consensus across scales.
3. **Consistency & Hierarchy Analysis**: We determine the token transition matrix to reveal the multi-scale hierarchy via computing the token similarity. The transition matrix could be used to translate multi-scale fold language without rerun the model.

### 2.2 Invariant Graph Encoder

Due to the rotation and translation equivariant nature, the same protein may have different coordinate records, posing a challenge in learning compact invariant representations. Previous works [4, 12, 2, 8] have shown that the invariant featurizer can extract informative structure patterns, and we follow the same road: representing the protein structures as a graph consisting of invariant node and edge features. We then use BlockGAT [8] to learn high-level representations.

#### Protein Block Graph

Given a protein 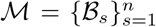 containing *n* blocks, where each block, represents an amino acid, we build the block graph 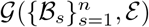using kNN algorithm. In the block graph, the *s*-th node is represented as ℬ_*s*_ = (*T*_*s*_, ***f***_*s*_), and the edge between (*s, t*) is represented as ℬ_*st*_ = (*T*_*st*_, ***f***_*st*_). *T*_*s*_ = (*R*_*s*_, ***t***_*s*_) and 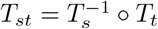 are the local frames of the *s*-th and the relative transform between the *s*-th and *t*-th blocks, respectively. ***f***_*s*_ and ***f***_*st*_ are the node and edge features. *R*_*s*_ and ***t***_*s*_ are the rotation and translation of the *s*-th block, serving as the local frame parameters.

#### BlockGAT Encoder

We use the BlockGAT [8] layer *f*_*θ*_ to learn block-level representations:

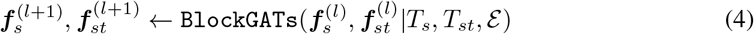

where 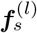 and 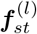 represent the input node and edge features of the *l*-th layer. *T*_*s*_ = (*R*_*s*_, ***t***_***s***_) is the local frame of the *s*-th block, and 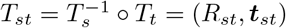 is the relative transform between the *s*-th and *t*-th blocks.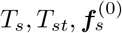 and 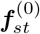 are initialized from the ground truth structures using the invariant featurizer proposed in UniIF [8]. We write the encoder’s output as **F** = [***f***_1_, ***f***_2_, ⋯, ***f***_*n*_], where ***f***_*s*_ is the embedding of the *s*-th residue, and refer to the encoding space as the semantic space.

### 2.3 Multiscale Code Generator

#### Multiscale Code

We introduce masked-binary code to serve as unified representation across scales. When the codebook size is 2^*m*^ and padding length *M*, the binary code of the decimal integer *z* is:

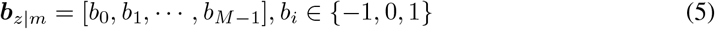

where *b*_*i*_ is the *i*-th bit of the binary code. Note that *b*_*i*_ = −1 if *i* < *M* − *m*; otherwise, *b*_(*M*−*m*):*M*−1_ is the binary form of *z*. For example, if *M* = 4, we have ***b***_0|2_ = [−1, −1, 0, 0], ***b***_1|2_ = [−1, −1, 0, 1], ***b***_3|2_ = [−1, −1, 1, 1], ***b***_7|3_ = [−1, 1, 1, 1], ⋯, ***b***_15|4_ = [1, 1, 1, 1]

#### Code Embedding via Adapter

A MLP, i.e., `CodeAdapter`_m_ : ℝ^M^ → ℝ^d^, is used to project ***b***_*i*|*m*_ ∈ ℝ^M^ to code vectors ***v***_*i*_ ∈ ℝ_d_ to consider correlations in each bit position. Formally, we write

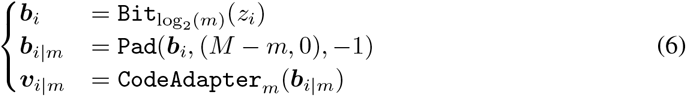

where `Bit`(·) convert decimal numbers to its binary form, ***b***_*i*|*m*_ is the masked binary vector, and ***v***_*j*_ is the *j*-th code embedding at scale *m*. If *m* = 10, the codebook size is 2^10^, and the MLP projects 1024 binary vectors into 1024 *d*-dimension code embeddings.

#### Code Consistency & Hierarchy

By training multiscale models simultaneously, the `CodeAdapter`_m_ projects the masked-binary code into the same semantic space, aligning with the output of the shared encoder. Therefore, one can discover the code mapping relationship between different scales by computing the similarity in the semantic space. For scale *m* and *m*′, where *m* > *m*′, we define a transition matrix **M**_*m*→*m*_*′* to record the code transition relationship:

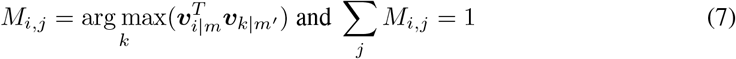

If *M*_*i,j*_ = 1, the *i*-th code at scale *m* is associate to the *j*-th code at scale *m*′. The transition matrix could be used to translate the code embeddings across scales without rerunning the model.

### 2.4 Quantifier: Code Alignment, Optimization, and Mixing

As in FoldToken3 [3], we apply a novel quantifier, called Binary Stochastic Quantifier (BSQ), to quantize the embeddings. The key problem is: *how to replace latent embedding* ***f***_*i*_ *with the most similar token embedding* ***v***_*j*_ *in a differential way?*

#### Find Neighbor

In vanilla vector quantization, they use “argmax” operation to find the nearest code vector, which is non-differential. In this paper, we take the selection process as sampling from a multi-class distribution *z*_*i*|*m*_ ∼ `Mult`(***p***_*i*|*m*_) at scale *m*:

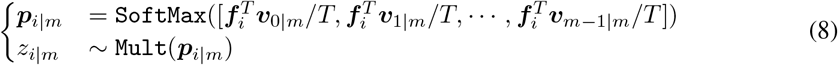

and then, we optimize ***p***_*i m*_ in a differential way. The temperature parameter *T* controls the softness of the attention query operation. When *T* is large, the attention weights will be close to uniform; otherwise, the attention weights will be close to one-hot.

#### Optimize Neighbor

In Eq. 8, the sampling operation is non-differentiable, and we use a reparameterization trick to optimize ***p***_*i*|*m*_:

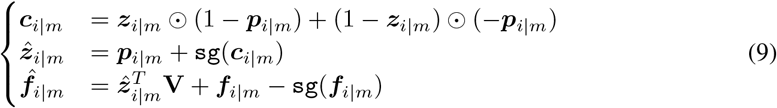

where `sg`(·) is the stop gradient operation, ⊙ is the element-wise multiplication, and ***z***_*i*|*m*_ ∈ ℝ_m_ is the onehot version of *z*_*i*_ at scale *m*. The first two equations reparameterize ***z***_*i*|*m*_ as 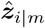 to allow ***p***_*i*|*m*_ get gradient, inspired by [10]; the third equation allows ***h***_*i*|*m*_ to get direct gradient for optimizing the encoder. 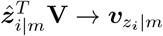 operation selects the *z*_*i*_-th code vector using the onehot 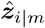.

#### Multiscale Code Mixing

During training, we uniformly sample the scale *m* and generate the code embeddings 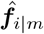 to serve as 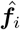:

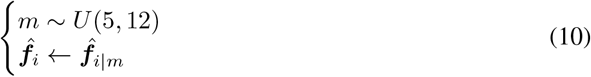

where *U* (*a, b*) is the uniform distribution between *a* and *b*. The average code embedding is a mixture of codes across scales:

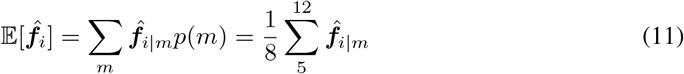

#### Teacher Guidance

To accelerate training convergence, we randomly copy encoder output ***f***_*i*_ as decoder input 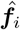 in probability *T* :

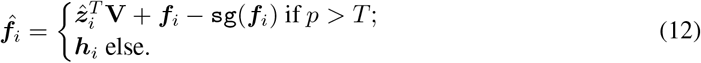

where *p* ∼ *U* (0, 1) is sampled from uniform distribution. When *T* = 1.0, the vector quantization module is skipped, allowing the encoder-decoder to be easily optimized. When *T* = 0.0, the vector quantization module is fully used. For values of 0.0 < *T* < 1.0, the shortcut guides the vector quantization model to learn code vectors aligning with the encoder’s inputs. The adaptive temperature scheduler is dependent on the loss *ℒ*:

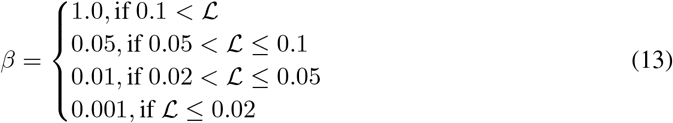

#### Stable Gradient

The scaled softmax operation in Eq. 9 bridges the continual model (*T* > 0) to discrete vector quantization (*T* = 0); thus allowing precise gradient computation rather than gradient mismatch in the VVQ. During training, we gradually anneal the temperature from 1.0 to 1e-8; however, the gradient of the scaled softmax tend to explode when *T* is extremely small:

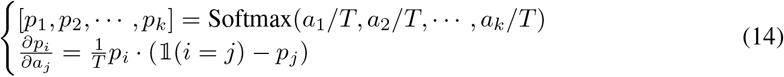

The unstable gradient would lead to representation and codebook collapses, as the `CodeAdapter`_m_ and encoder parameters collapse after one step of updating a large gradient. To overcome the issue, we introduce the trick of ‘partial gradient’:

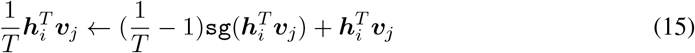

where the first term is stoped gradient and only the second term contribute to gradient computation. Obviously, Eq.15 has the same forward behavior like Eq.9 while the gradient is stable and do not affected by the extreme small value of *T*.

### 2.5 Equivariant Graph Decoder

Generating the protein structures conditioned on invariant representations poses significant challenges in computing efficiency. For example, training well-known AlphaFold2 from scratch takes 128 TPUv3 cores for 11 days [13]; OpenFold takes 50000 GPU hours for training [1]. In this work, we propose an efficiency plug-and-play SE(3)-layer that could be added to any GNN layer for structure prediction. Thanks to the simplified module of the SE(3)-layerand BlockGAT with sparse graph attention, we can train the model on the entire PDB dataset in 1 day using 8 NVIDIA-A100s.

#### SE-(3) Frame Passing Layer

We introduce frame-level message passing, which updates the local frame of the *s*-th block by aggregating the relative rotation *R*_*s*_ and translation ***t***_*s*_ from its neighbors:

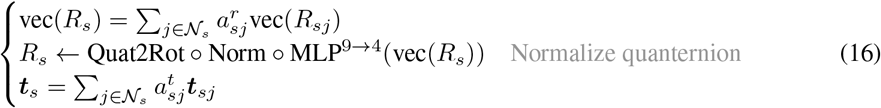

where 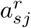 and 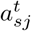 are the rotation and translation weights, and *𝒩*_*s*_ is the neighbors of the *s*-th block. vec(·) flattens 3 × 3 matrix to 9-dimensional vector. MLP^9→4^(·) maps the 9-dim rotation matrix to 4-dim quaternion, and Norm(·) normalize the quaternion to ensure it represents a valid rotation. Quat2Rot(·) is the quaternion to rotation function. We further introduce the details as follows:

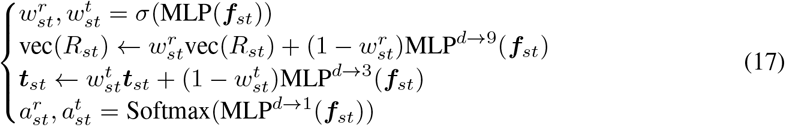

where 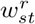 and 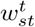 are the updating weights for rotation and translation, 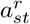 and 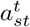 are the attention weights. The proposed SE-(3) layer could be add to any GNN for local frame updating.

#### Iterative Refinement

We propose a new module named SE-(3) BlockGAT by adding a SE-(3) layer to BlockGAT. We stack multi-layer SE-(3) BlockGAT to iteratively refine the structures:

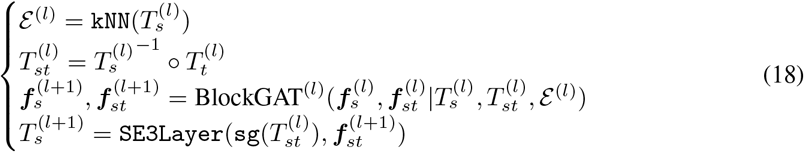

where `sg`(·) is the stop-gradient operation, and `SE3Layer`(·) is the SE-(3) layer described in Eq.17. Given the predicted local frame 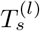, we can obtain the 3D coordinates by:

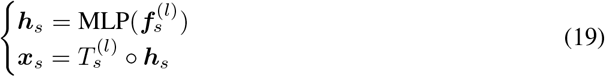

### 2.6 Reconstruction Loss

Inspired by Chroma [11], we use multiple loss functions to train the model. The overall loss is:

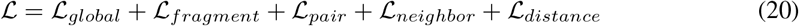

To illustrate the loss terms, we define the aligned RMSD loss as 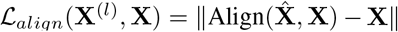, given the the ground truth 3D coordinates **X** ∈ ℝ^*n*,3^ and the predicted 3D coordinates 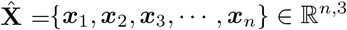. The global, fragment and pair loss are defined by the aligned MSE loss, but with different input data shape:

- **Global Loss**: **X** with shape [*n*, 4, 3]. RMSD of the global structure.
- **Fragment Loss**: **X** with shape [*n, c*, 4, 3]. RMSD of *c* neighbors for each residue.
- **Pair Loss**: **X** with shape [*n, K, c* · 2, 4, 3]. RMSD of *c* neighbors for each kNN pair.
- **Neighbor Loss**: **X** with shape [*n, K*, 4, 3]. RMSD of *K* neighbors for each residue.

where *n* is the number of residues, *c* = 7 is the number of fragments, *K* = 30 is the number of kNN, 4 means we consider four backbone atoms {*N, CA, C, O*}, and 3 means the 3D coordinates. The distance loss is defined as the MSE loss between the predicted and ground truth pairwise distances:

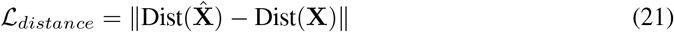

where Dist(**X**) ∈ ℝ^*n,n*^ is the pairwise distance matrix of the 3D coordinates **X**. We apply the loss on each decoder layer, and the final loss is the average, whcih is crucial for good performance.

## 3 Experiments

We conduct systematic experiments to evaluate FoldToken4:

- **Single-Chain Benchmark (Q1)**: How well FoldToken4 perform on single-chain data?
- **Multi-Chain Benchmark (Q2)**: How well FoldToken4 perform on multi-chain data?
- **Code Analysis (Q3):** What can we learn from FoldToken4’s hierarchical code space?

### Multi-chain PDB Data for Training

We train the model using all proteins collected from the PDB dataset as of 1 March 2024. After filtering residues with missing coordinates and proteins less than 30 residues, we obtain 162K proteins for training. We random crop proteins to ensure that the length range from 5 to 1024. Protein complexes are supported by adding chain encoding features *c*_*ij*_ to the edge *e*_*ij*_: *c*_*ij*_ = 0, if *i* and *j* are in different chains; else *c*_*ij*_ = 1. The model is trained for up to 60 epochs with a batch size of 8 and a learning rate of 0.001, using 8 NVIDIA-A100s.

### Metrics

Regarding reconstruction, we evaluate the model using the average TMscore and aligned RMSD. In FoldToken2, we uses Kabsch algorithm to align the predicted structure to the ground truth structure; however, the aligned RMSD seems to be different to that of PyMol. We do not know what is the reason for this discrepancy. Finally, we use PyMol’s API for computing RMSD. We also introduce a similarity metric to evaluate the codebook diversity:

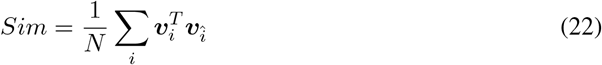

where ***v***_*i*_ and ***v***_î_ are the *i*-th and nearest neighbor code vectors, respectively. The similarity metric ranges from -1 to 1, with 1 indicating that there is always a very similar code vector for each one. To make discrete tokens distinguishable, the smaller the similarity, the better the diversity, and the easier it is to predict. Considering the extreme case where the similarity is 1, one code id can be replaced by that of its nearest neighbor without affecting the reconstruction quality, leading to inconsistent fold languages. Also, high similarity indicates that the model do not robust to noise of the embeddings, as similar code vectors may be easily confused by each other.

### 3.1 Single-Chain Benchmark (Q1)

#### Single-Chain Data for Evaluation

Following FoldToken2, we evaluate models on T493 and T116 datasets for head-to-head comparison, which contains 493 and 116 proteins, respectively. We also evaluate methods on 128 novel proteins released after the publication of AlphaFold3, called N128. In Table. 1, we show the reconstruction results of FoldToken4 and conclude that:

**Table 1:** Single-chain Reconstruction Benchmark. FT1, FT2, FT3, and FT4 indicates FoldToken1 [5, 7], FoldToken2 [6], FoldToken3 [3], and FoldToken4, respectively. We also report the reconstruction results of ESM3 [9] for comprehensive understanding. When KNN is ‘full’, the approach uses full attention.

| Model | Data | Config | | | | | TMScore ↑ | | | RMSD ↓ | | |
| --- | --- | --- | --- | --- | --- | --- | --- | --- | --- | --- | --- | --- |
|  |  | #Code | #Enc | #Dec | #Hid | #KNN | Avg | Max | Min | Avg | Max | Min |
| FT1 | T116 | 65536 | 12 | 12 | 480 | full | 0.77 | 0.96 | 0.39 | 3.31 | 24.53 | 0.52 |
| FT2 | T116 | 65536 | 8 | 8 | 128 | 30 | 0.97 | 0.99 | 0.90 | 0.52 | 0.76 | 0.30 |
| ESM3 | T116 | 4096 | 2 | 30 | 1024 | full | 0.97 | 0.99 | 0.76 | 1.97 | 13.27 | 0.01 |
| FT3 | T116 | 4096 | 8 | 8 | 128 | 30 | 0.95 | 0.98 | 0.88 | 0.64 | 0.98 | 0.30 |
| FT3 | T116 | 1024 | 8 | 8 | 128 | 30 | 0.93 | 0.97 | 0.83 | 0.73 | 1.20 | 0.26 |
| FT3 | T116 | 256 | 8 | 8 | 128 | 30 | 0.93 | 0.98 | 0.86 | 0.76 | 1.10 | 0.44 |
| FT3 | T116 | 128 | 8 | 8 | 128 | 30 | 0.91 | 0.96 | 0.82 | 0.87 | 1.36 | 0.46 |
| FT3 | T116 | 64 | 8 | 8 | 128 | 30 | 0.89 | 0.96 | 0.74 | 1.02 | 1.80 | 0.32 |
| FT4 | T116 | 4096 | 8 | 8 | 128 | 30 | 0.95 | 0.98 | 0.78 | 0.67 | 1.25 | 0.28 |
| FT4 | T116 | 2048 | 8 | 8 | 128 | 30 | 0.94 | 0.98 | 0.82 | 0.70 | 1.08 | 0.27 |
| FT4 | T116 | 1024 | 8 | 8 | 128 | 30 | 0.94 | 0.98 | 0.79 | 0.71 | 1.11 | 0.30 |
| FT4 | T116 | 512 | 8 | 8 | 128 | 30 | 0.94 | 0.98 | 0.81 | 0.73 | 1.19 | 0.27 |
| FT4 | T116 | 256 | 8 | 8 | 128 | 30 | 0.93 | 0.98 | 0.80 | 0.78 | 1.26 | 0.30 |
| FT4 | T116 | 128 | 8 | 8 | 128 | 30 | 0.92 | 0.97 | 0.79 | 0.86 | 1.54 | 0.28 |
| FT4 | T116 | 64 | 8 | 8 | 128 | 30 | 0.89 | 0.96 | 0.73 | 1.02 | 1.97 | 0.29 |
| FT4 | T116 | 32 | 8 | 8 | 128 | 30 | 0.86 | 0.94 | 0.62 | 1.22 | 2.12 | 0.31 |
| FT1 | T493 | 65536 | 12 | 12 | 480 | full | 0.74 | 0.96 | 0.44 | 3.09 | 18.09 | 0.48 |
| FT2 | T493 | 65536 | 8 | 8 | 128 | 30 | 0.95 | 0.99 | 0.78 | 0.64 | 1.99 | 0.36 |
| ESM3 | T493 | 4096 | 2 | 30 | 1024 | full | 0.95 | 0.99 | 0.32 | 2.40 | 15.97 | 0.01 |
| FT3 | T493 | 4096 | 8 | 8 | 128 | 30 | 0.92 | 0.99 | 0.57 | 0.86 | 6.48 | 0.35 |
| FT3 | T493 | 1024 | 8 | 8 | 128 | 30 | 0.90 | 0.98 | 0.61 | 0.98 | 7.52 | 0.38 |
| FT3 | T493 | 256 | 8 | 8 | 128 | 30 | 0.90 | 0.98 | 0.52 | 1.03 | 6.14 | 0.38 |
| FT3 | T493 | 128 | 8 | 8 | 128 | 30 | 0.88 | 0.97 | 0.49 | 1.15 | 7.70 | 0.39 |
| FT3 | T493 | 64 | 8 | 8 | 128 | 30 | 0.85 | 0.96 | 0.45 | 1.33 | 7.47 | 0.47 |
| FT4 | T493 | 4096 | 8 | 8 | 128 | 30 | 0.92 | 0.98 | 0.55 | 0.95 | 5.88 | 0.33 |
| FT4 | T493 | 2048 | 8 | 8 | 128 | 30 | 0.91 | 0.98 | 0.59 | 0.95 | 6.00 | 0.37 |
| FT4 | T493 | 1024 | 8 | 8 | 128 | 30 | 0.91 | 0.98 | 0.54 | 0.98 | 7.18 | 0.31 |
| FT4 | T493 | 512 | 8 | 8 | 128 | 30 | 0.91 | 0.98 | 0.56 | 1.02 | 5.78 | 0.36 |
| FT4 | T493 | 256 | 8 | 8 | 128 | 30 | 0.90 | 0.97 | 0.58 | 1.07 | 6.90 | 0.41 |
| FT4 | T493 | 128 | 8 | 8 | 128 | 30 | 0.88 | 0.97 | 0.56 | 1.17 | 6.75 | 0.39 |
| FT4 | T493 | 64 | 8 | 8 | 128 | 30 | 0.84 | 0.95 | 0.47 | 1.40 | 7.98 | 0.36 |
| FT4 | T493 | 32 | 8 | 8 | 128 | 30 | 0.80 | 0.94 | 0.44 | 1.71 | 7.77 | 0.48 |
| FT1 | N128 | 65536 | 12 | 12 | 480 | full | 0.62 | 0.93 | 0.26 | 11.20 | 53.25 | 0.47 |
| FT2 | N128 | 65536 | 8 | 8 | 128 | 30 | 0.94 | 0.99 | 0.17 | 0.78 | 5.88 | 0.27 |
| ESM3 | N128 | 4096 | 2 | 30 | 1024 | full | 0.92 | 1.00 | 0.30 | 4.50 | 22.51 | 0.04 |
| FT3 | N128 | 4096 | 8 | 8 | 128 | 30 | 0.92 | 0.99 | 0.26 | 1.16 | 4.42 | 0.41 |
| FT3 | N128 | 1024 | 8 | 8 | 128 | 30 | 0.91 | 0.98 | 0.29 | 1.25 | 4.31 | 0.39 |
| FT3 | N128 | 256 | 8 | 8 | 128 | 30 | 0.89 | 0.98 | 0.22 | 1.49 | 7.76 | 0.37 |
| FT3 | N128 | 128 | 8 | 8 | 128 | 30 | 0.89 | 0.96 | 0.27 | 1.60 | 6.78 | 0.39 |
| FT3 | N128 | 64 | 8 | 8 | 128 | 30 | 0.84 | 0.95 | 0.23 | 2.34 | 15.64 | 0.49 |
| FT4 | N128 | 4096 | 8 | 8 | 128 | 30 | 0.91 | 0.99 | 0.21 | 1.34 | 7.75 | 0.38 |
| FT4 | N128 | 2048 | 8 | 8 | 128 | 30 | 0.91 | 0.98 | 0.21 | 1.37 | 8.96 | 0.38 |
| FT4 | N128 | 1024 | 8 | 8 | 128 | 30 | 0.90 | 0.98 | 0.22 | 1.42 | 8.87 | 0.46 |
| FT4 | N128 | 512 | 8 | 8 | 128 | 30 | 0.90 | 0.98 | 0.21 | 1.43 | 8.39 | 0.53 |
| FT4 | N128 | 256 | 8 | 8 | 128 | 30 | 0.89 | 0.98 | 0.21 | 1.53 | 7.37 | 0.33 |
| FT4 | N128 | 128 | 8 | 8 | 128 | 30 | 0.88 | 0.97 | 0.20 | 1.64 | 9.67 | 0.52 |
| FT4 | N128 | 64 | 8 | 8 | 128 | 30 | 0.85 | 0.95 | 0.20 | 2.02 | 8.64 | 0.65 |
| FT4 | N128 | 32 | 8 | 8 | 128 | 30 | 0.78 | 0.93 | 0.11 | 2.91 | 12.76 | 0.86 |

#### FoldToken4 is comparable to FoldToken3 using a unified model

Although FoldToken4 is trained once for all scales, it achieves comparable reconstruction performance to FoldToken3, which is trained separately for each scale. The phenomenon is consistent across multiple datasets, including T493, T116, and N128, and multiple scales, including 2^6^, 2^7^, · · ·, 2^12^. We emphasize that it is non-trival to jointly learn multi-scale models at different scales without performance drop, as the model has to strike a balance between different scales, like mult-task learning. The keys to the good performance are the proposed conditional code adapters and token mixing operation.

#### FoldToken4 supports minimum 32 code size

Beyond FoldToken3, FoldToken4 can support a minimum code size of 32, which is the lowest code size feasible in our experiments. If we futher reduce the code size to 16, the model could not reconstruct reasonable protein structures. From 2^5^ to 2^12^ code sizes, we find that the model achieves performance plateau in the code size of 128. Therefore, considering both reconstruction quality and compression ratio, we recommend using a code size of 128 or 256 for FoldToken4. However, smaller codebook like 64 or 32 can be used for structure alignment or similarity search tasks, as the algorithm do not require exact structure details. In Fig. 2, we show the cases with codebook size 32 in N128. We observe that for proteins with fewer than 500 amino acids, the model can provide reasonable reconstruction results. When the protein is too long, the model may fail to recover fine secondary structures, but the coarse global shape is well preserved. It is amazing to find that the 32 words can reconstruct the protein structures effectively.

**Figure 2:**
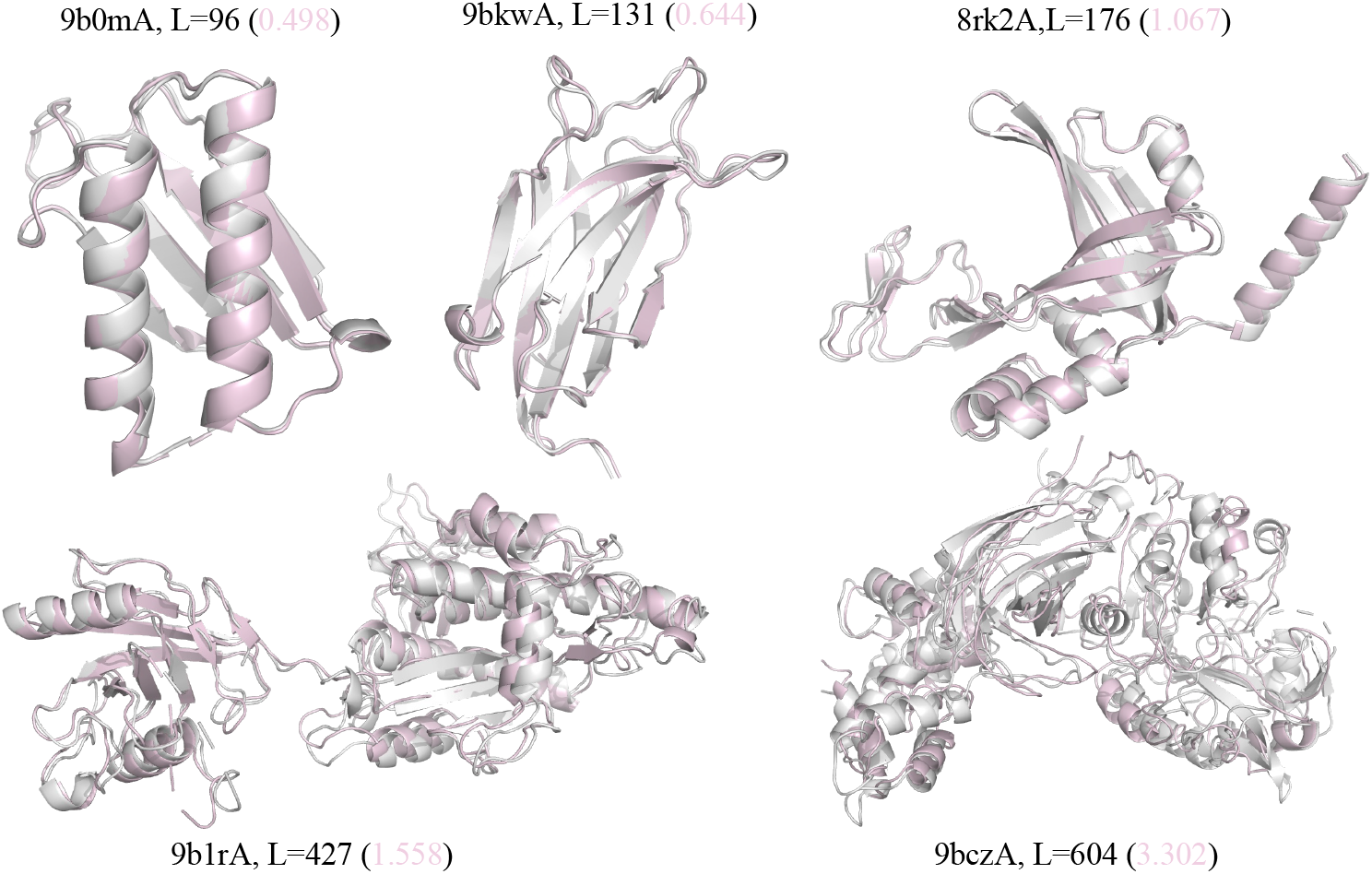
The cases with codebook size 32 in N128. Grey structures are the ground truth, and colored structures are the reconstructed ones by FoldToken3 and FoldToken4. We provide the RMSD in the brackets.

#### When FoldToken4 will fail in single-chain data?

In Fig. 3, we show cases in N128 where the TMScore is less than 0.5, the the codebook size of 256. We observe that both FoldToken3 and FoldToken4 model have low TMScore when the protein is too short. This is because proteins with fewer than 30 amino acids were filtered out in the training set. Nevertheless, we can see that the global shape of the protein is still well preserved, although the accuracy of the secondary structure is not satisfactory. Moreover, FoldToken4 generally performs better than FoldToken3 in these cases.

**Figure 3:**
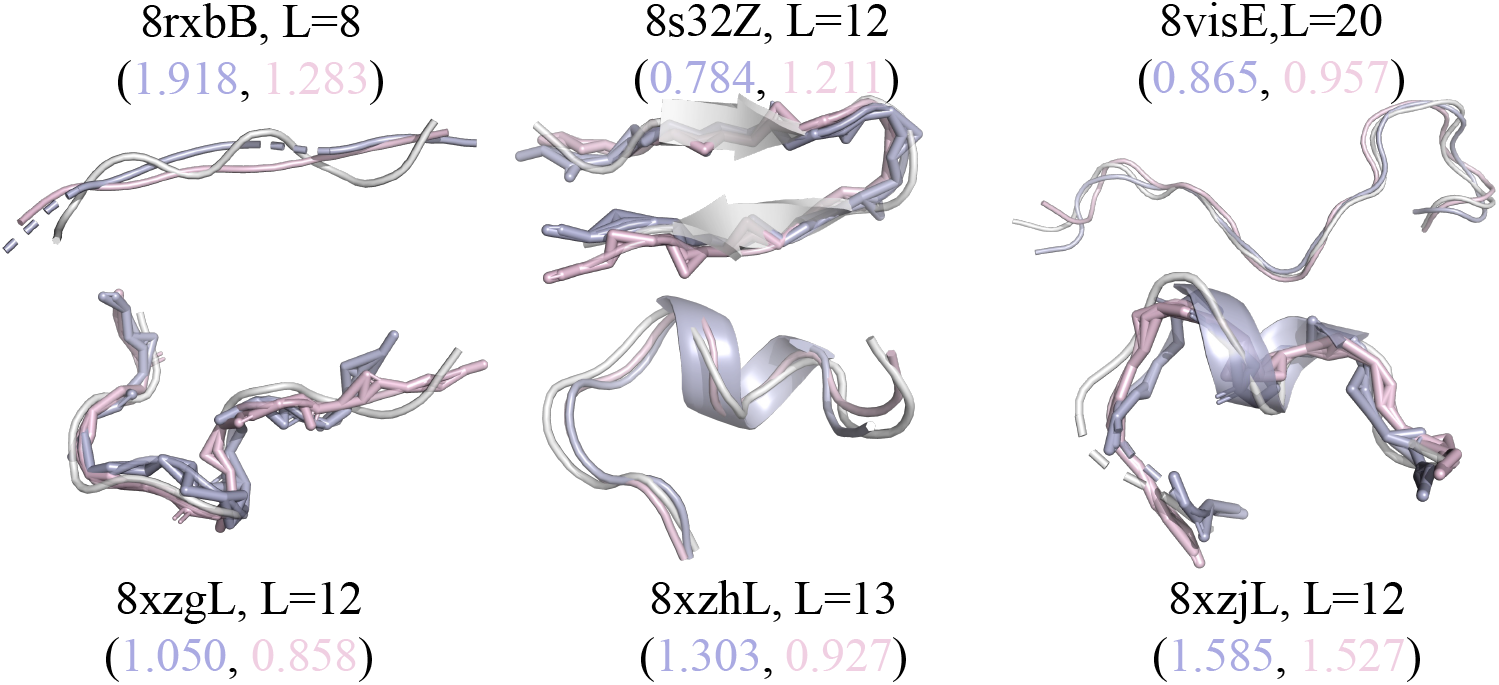
The cases when TMScore<0.5 in N128. Grey structures are the ground truth, and colored structures are the reconstructed ones by FoldToken3 and FoldToken4. We provide the RMSD in the brackets.

#### FT4 is comparable to ESM3 using less parameters and data

When comparing FoldToken4 with ESM3, we find that FoldToken4 achieves comparable reconstruction performance to ESM3 while using fewer parameters and less data. In terms of trainable parameters, the encoder and decoder of FoldToken4 have **4.31M and 4.92M** parameters, respectively. In comparison, ESM3’s encoder and decoder have **30.1M and 618.6M** parameters, respectively. Regarding the training data, FoldToken4 is trained on the PDB dataset, which is a small subset of ESM3’s training set. Additionally, FoldToken4 is specifically trained for multi-chain protein reconstruction, a more challenging task than the single-chain protein reconstruction that ESM3 is trained for. Nevertheless, the checkpoints learned from the multi-chain task generalize well to single-chain tasks.

### 3.2 Multi-Chain Benchmark (Q2)

#### Multi-Chain Data for Evaluation

We evaluate the model on the antibody-antigen dataset (SAbDab), which contains 6741 protein complexes. We use foldseek to cluster protein chains into 1323 clusters:

~~~
foldseek easy - cluster ab_pdb / res tmp -c 0.7
~~~

The representative chains of each cluster are used for evaluation. After filtering representative proteins with less than 30 residues or more than 1000 residues, we get the evaluation dataset containing 1031 protein complexes, named M1031.

In Table. 2, we show the reconstruction results of FoldToken4 and conclude that:

**Table 2:**
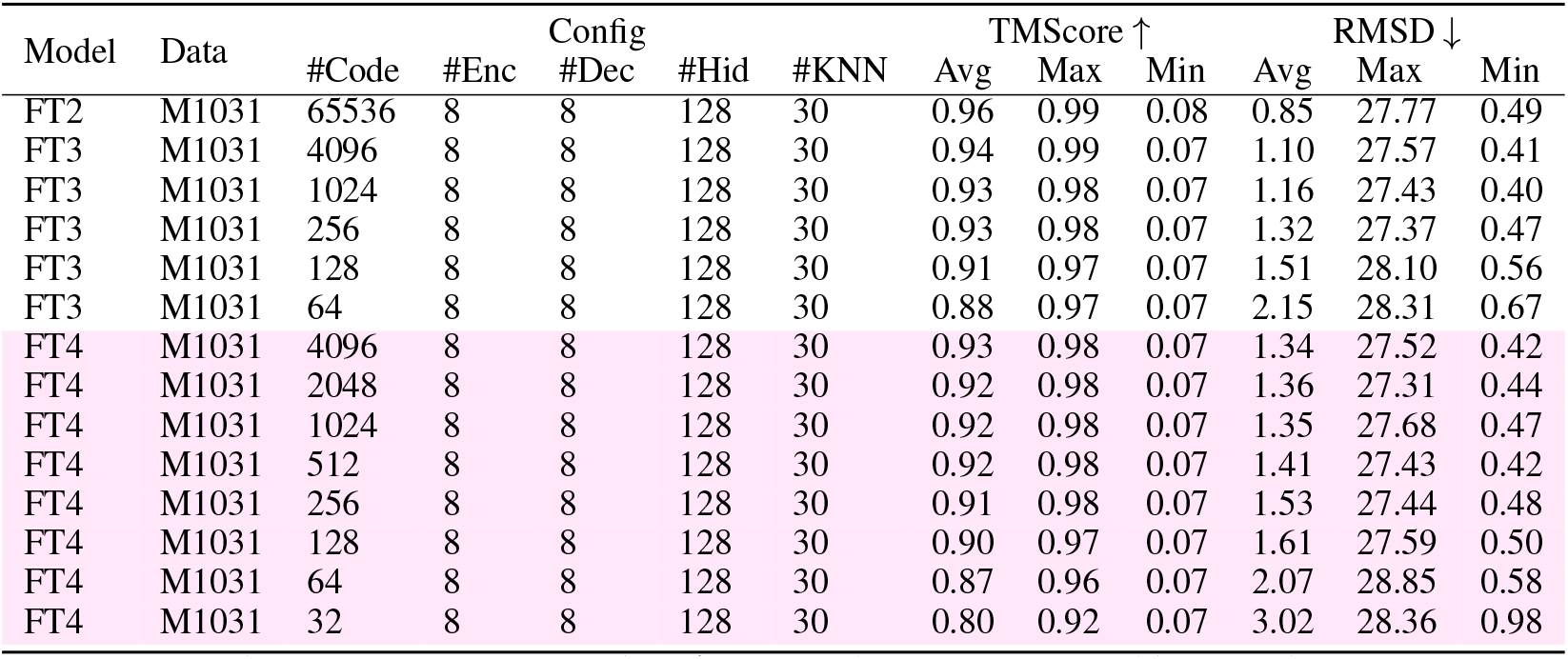
Multi-chain Reconstruction Benchmark. FT1 and ESM3 are ignored here, as they cannot process multi-chain proteins.

#### The compact code space can represent complex interactions

When using 256 code vectors or less, FoldToken4 can reconstruct protein complex structures effectively. Before FoldToken2, we believed that tokenizing single-chain proteins was difficult, especially with very small codebooks. However, FoldToken4 demonstrates that even protein complexes can be represented well using a small codebook, which is indeed surprising. This discovery will promote complex modeling, such as similar interface searching, complex alignment, and complex generation.

#### Learning multi-scale consistent models slightly reduces the reconstruction performance

In Table. 2, we observe that FoldToken4 achieves slightly worse performance than FoldToken3 with the same codebook size. This is because the model has to learn multi-scale tokenization in the same training session, where the learning objectives may conflict with each other. Considering the slight performance drop is acceptable, while the model can provide a consistent fold language across scales, we believe FoldToken4 would have more application scenarios. In Fig.4, we show some the largest reconstructed protein systems using FoldToken4 with codebook size of 4096.

**Figure 4:**
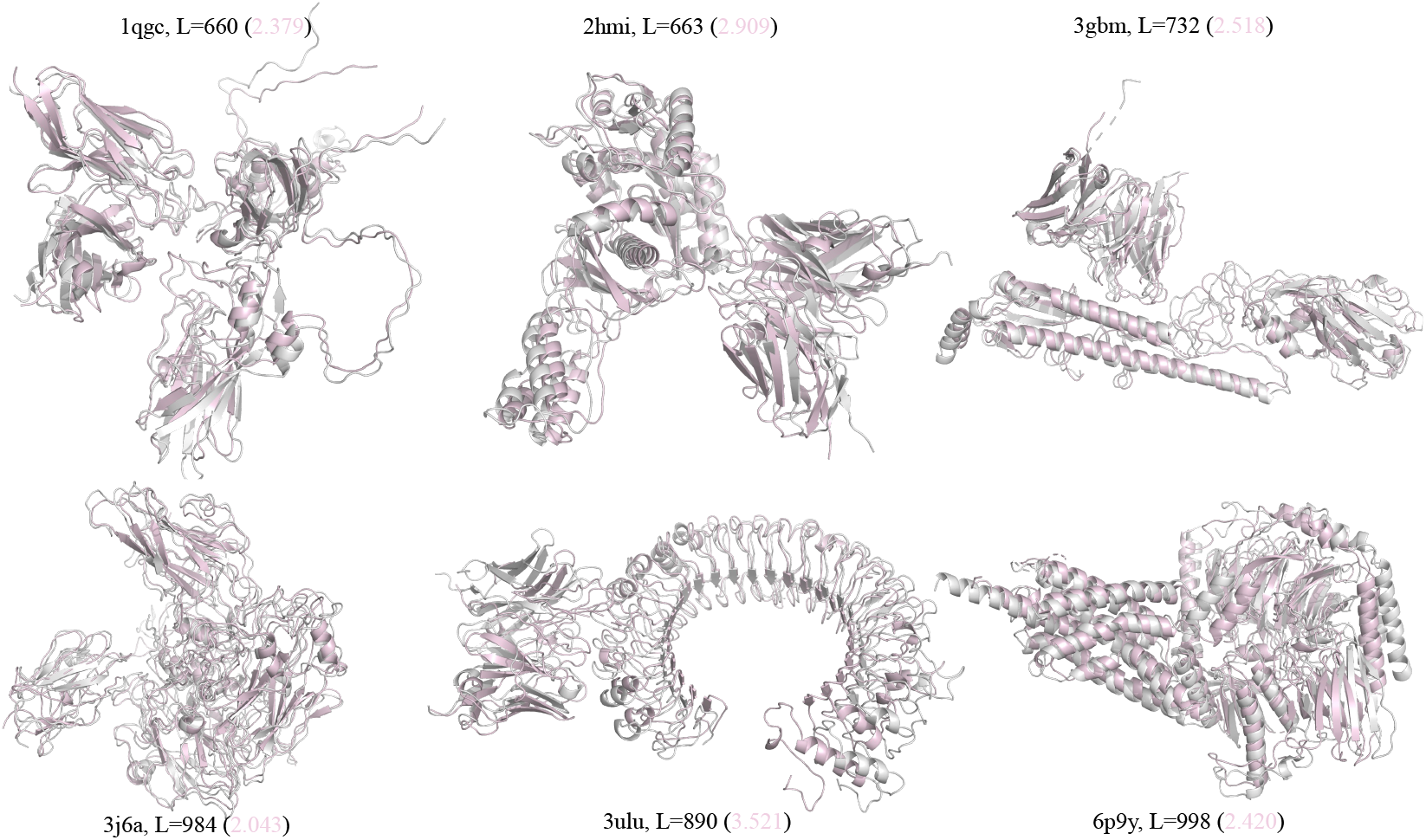
Large protein complexes reconstructed by FoldToken4 with codebook size of 4096. Grey structures are the ground truth, and colored structures are the reconstructed ones. We provide the RMSD in the brackets.

## 4 Consistency and Hierarchy (Q3)

### Code Diversity

We compute the cosine similarity for each code and its nearest neighbor and show the code similarity; refer to Eq. 22. In Table. 3, we observe that FoldToken4 has the most diverse code vectors, indicating the learnged language is more distinguishable than FoldToken1&2&3. In addition, the code diversity increases as the codebook size decreases, which is consistent with our expectation: the `CodeAdapter` should learn uniformly distributed code vectors in the encoding space to represent the protein structures effectively.

**Table 3:** The code vector similarity.

| Model | #Code | Avg↓ | Max↓ | Min↓ |
| --- | --- | --- | --- | --- |
| FT1 | 65536 | 0.9959 | 1.0 | 0.7447 |
| FT2 | 65536 | 0.9999 | 1.0 | 0.9625 |
| FT3 | 256 | 0.9070 | 0.9711 | 0.7371 |
| FT4 | 4096 | 0.9872 | 0.9999 | 0.7563 |
| FT4 | 2048 | 0.9776 | 0.9995 | 0.8045 |
| FT4 | 1024 | 0.9602 | 0.9991 | 0.7673 |
| FT4 | 512 | 0.9346 | 0.9940 | 0.7949 |
| FT4 | 256 | 0.8962 | 0.9693 | 0.7294 |
| FT4 | 128 | 0.8389 | 0.9064 | 0.7047 |
| FT4 | 64 | 0.7775 | 0.8551 | 0.6763 |
| FT4 | 32 | 0.7098 | 0.7956 | 0.6113 |

### Consistency

FoldToken4 can generate consistent fold languages across scales, as shown in Fig. 5. The larger the code size, the more detailed structures will be preserved. Thanks to the shared encoder-decoder architecture, multi-scale code vectors can be projected to the same space, and the hierarchical token-mapping relationships across scales can be discovered; refer to Fig. 6. One can also fuse multi-scale languages, where complex structures use more detailed tokens, and simple structures use fewer tokens.

**Figure 5:**
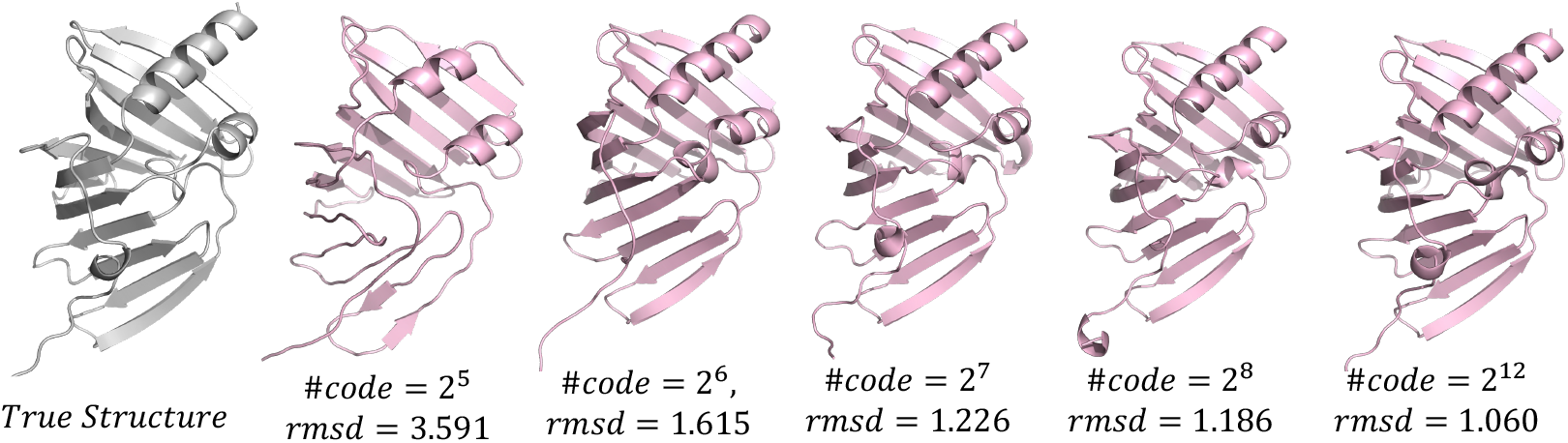
Multi-scale structures generated by FoldToken4.

**Figure 6:**
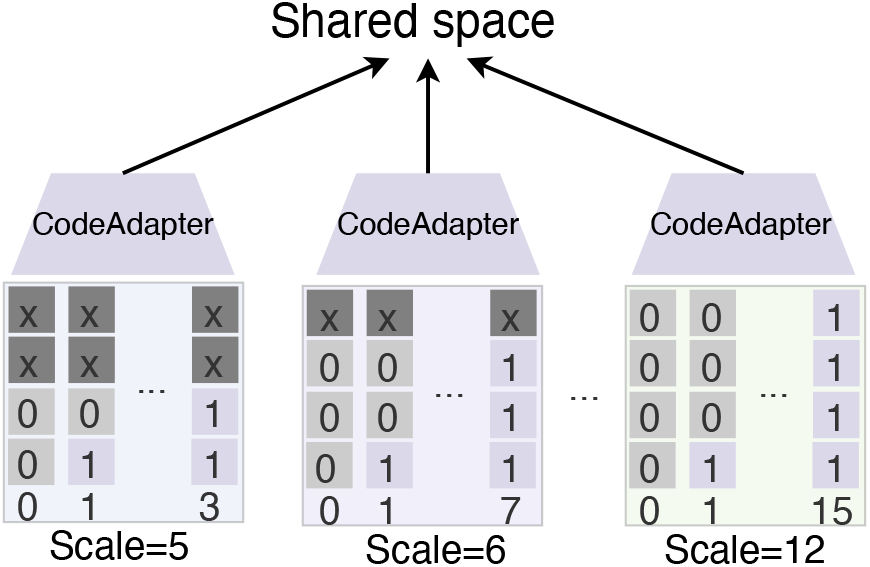
Multi-scale models share the same encoding space.

### Hierarchy

In the shared encoding space, we compute the token similarity across scales to get the token-mapping matrices, as illustrated by Eq. 7. Fig. 7 shows examples of hierarchically translating fold languages from scale=12 to scale=5 using the learned token-mapping matrices. We observe that, without rerunning the model, we can generate multi-scale fold languages from the most detailed scale to the coarsest scale with good reconstruction quality. This approach is highly efficient in both computation and storage: one only needs to record languages at scale 12, and can then generate languages at other scales using the token-mapping matrices.

**Figure 7:**
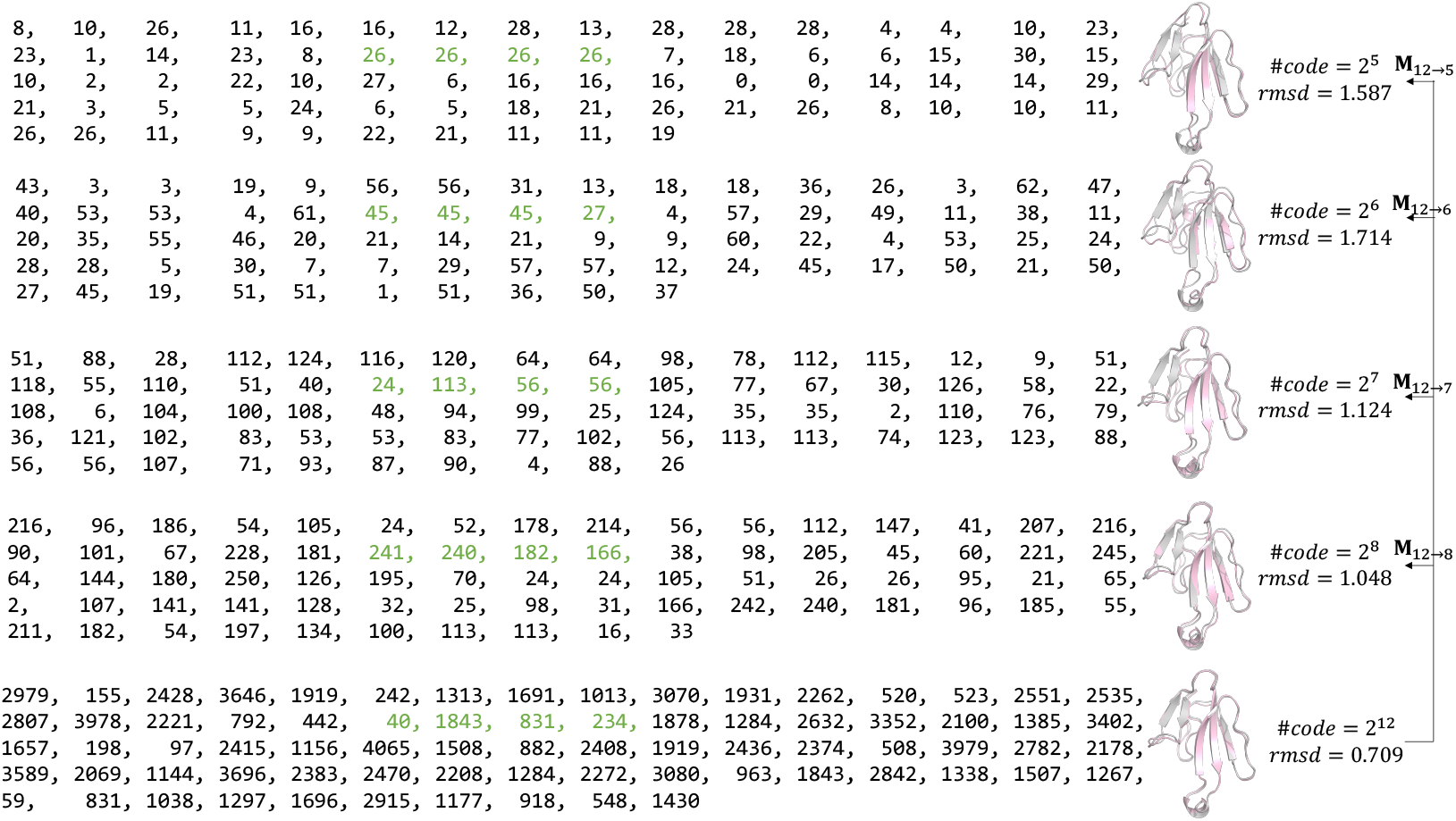
Hierarchical language transition from fine to coarse. We generate fold languages from the detailed scale (2^12^) to coarse scales using the token-mapping matrices, and visualize the reconstructed structures.

## 5 Conclusion

This paper introduces FoldToken4, a novel protein structure tokenization method that learns multi-scale consistent fold languages by introducing `CodeAdapter` and token-mixing techniques. Fold-Token4 offers advantages in consistency, hierarchy, and efficiency, in addition to FoldToken2&3’s invariant, compact, and generative merits. This advancement will benefit a wide range of protein structure-related tasks, including protein structure alignment, generation, and representation learning.

